# Correlation of *in vitro* biofilm formation capacity with persistence of antibiotic-resistant *Escherichia coli* on gnotobiotic lamb’s lettuce

**DOI:** 10.1101/2025.02.07.637020

**Authors:** Rudolf O. Schlechter, Elisabet Marti, Mitja N. P. Remus-Emsermann, David Drissner, Maria-Theresia Gekenidis

## Abstract

Bacterial contamination of fresh produce is a growing concern for food safety, as apart from human pathogens, antibiotic-resistant bacteria can persist on fresh leafy produce. A prominent persistence trait in bacteria is biofilm formation, as it provides increased tolerance to stressful conditions. We screened a comprehensive collection of 174 antibiotic-susceptible and -resistant *Escherichia coli* originating from fresh leafy produce and its production environment. We tested the ability of these strains to produce biofilms, ranging from none or weak to extreme biofilm-forming bacteria. Next, we tested the ability of selected antibiotic-resistant isolates to colonize gnotobiotic lamb’s lettuce (*Valerianella locusta*) plants. We hypothesized that a higher *in vitro* biofilm formation capacity correlates with increased colonization of gnotobiotic plant leaves. Despite a marked difference in the ability to form *in vitro* biofilms for a number of *E. coli* strains, *in vitro* biofilm formation was not associated with increased survival on gnotobiotic *V. locusta* leaf surfaces. However, all tested strains persisted for at least 21 days, highlighting potential food safety risks through unwanted ingestion of resistant bacteria. Population densities of biofilm-forming *E. coli* exhibited a complex pattern, with subpopulations more successful in colonizing gnotobiotic *V. locusta* leaves. These findings emphasise the complex behaviour of antibiotic-resistant bacteria on leaf surfaces and their implications to human safety.

**Importance:** Each raw food contains a collection of microorganisms, including bacteria. This is of special importance for fresh produce such as leafy salads or herbs, as these foods are usually consumed raw or after minimal processing, whereby higher loads of living bacteria are ingested than with a food that is heated before consumption. A common bacterial lifestyle involves living in large groups embedded in secreted protective substances. Such bacterial assemblies, so-called biofilms, confer high persistence and resistance of bacteria to external harsh conditions. In our research we investigated whether stronger *in vitro* biofilm formation of antibiotic-resistant *Escherichia coli* correlates with a better survival on lamb’s lettuce leaves. Although no clear correlation was observed between biofilm formation capacity and population density on the salad, all tested isolates could survive for at least three weeks with no significant decline over time, highlighting a potential food safety risk independently of *in vitro* biofilm formation.

## Introduction

Fresh produce is an important part of a healthy human diet, as regular consumption of salad and raw vegetables improves serum nutrient levels (1). However, fresh produce can be colonized by potentially harmful human pathogens that cause disease and foodborne outbreaks (2, 3). A recent meta-analysis describing fresh produce-related worldwide outbreaks from 1980 to 2016 revealed that fresh leafy produce accounted for more than half of the reported outbreaks (4), with almost 73,000 infections and 173 deaths associated to these outbreaks. The most important bacterial pathogens frequently associated with fresh produce outbreaks are *Salmonella* spp., certain strains of *Escherichia coli*, and *Listeria monocytogenes*. There are many possible contamination routes in the fresh produce production environment. The main sources of fresh produce contamination are irrigation water, soil, applied fertilizer, equipment, workers, or animals (3).

In contrast to pathogens which can cause direct illness and are therefore well-known through related foodborne outbreaks, antibiotic-resistant bacteria (ARB) which are frequently non-pathogenic commensals or environmental bacteria (5–7) often pass unnoticed despite their clinical relevance. Contamination events may spread pathogens as well as ARB on crops. Consequently, clinically-relevant antibiotic resistance genes (AGRs) have been widely detected on produce (8–14). This is further intensifying the global threat to health and food security as reported by the WHO (15). The WHO emphasizes the importance of hygienic production and processing of fresh produce, often consumed raw or minimally processed, to prevent ingestion of pathogens or ARB. Vegetables and minimally processed foods have been suggested as the main source of ARGs in the human gut microbiota (16). Human exposure to antibiotic-resistant (AR) *E. coli* strains through consumption of surface water-irrigated lettuce has been modelled to range between 10^−2^ and 10^6^ colony forming units (CFU) per 100 g of lettuce (17).

Pathogens and ARB must be able to grow or at least persist on edible plant parts after a contamination event to be transmitted from fresh produce to the consumer. The aboveground parts or plants, mainly represented by leaves, are collectively known as the phyllosphere, which offers an environment for microbes to grow (18). Although considered oligotrophic due to the heterogeneous and scarce availability of nutrients and water, bacteria are able to metabolise nutrients that are leached onto the leaf surface (19). Bacterial survival in the phyllosphere has been associated to aggregate sizes (20), suggesting that biofilm formation could be an important persistence factor.

Bacterial biofilms are surface-attached complex associations, embedded in a self-secreted matrix consisting mainly of polysaccharides, proteins, and extracellular DNA (21). The matrix protects bacterial cells from harsh conditions such as environmental stressors (e.g., UV radiation) (22), antibacterial agents, or the plant immune system (23, 24), potentially reducing the efficiency of sanitizing procedures. Commonly used sanitizing procedures applied in the food industry in several countries include various chlorinated agents, ozonated water, or UV radiation (25). Notably, despite the protective role of the biofilm matrix, plant chemical defences against colonization of bacterial pathogens have been proposed as promising alternatives to reduce bacterial biofilm formation in the phyllosphere (26). Additionally, the plant defence mechanism can be stimulated for more effective reduction of undesired bacteria colonizing fresh produce, for example by the use of UV-C radiation or ozone (27, 28).

Several models have been developed over decades to study biofilm formation *in vitro*, varying between continuous, semi-continuous, and batch culture (29). A well-established system suitable for high-throughput screenings is the crystal violet assay, which uses multiwell plates to test for bacterial production of biofilms after growth (29). Biofilm formation of antibiotic-resistant environmental strains on plant leaves has not been extensively studied, while biofilm formation of pathogenic strains has been well studied (reviewed by Yaron and Römling (26)). The review by Yaron and Römling (26) summarizes studies on biofilm formation by enteric pathogens such as *Salmonella enterica* or *E. coli* on produce surfaces and its role in survival on plants. For example, *S. enterica* survival has been assessed on parsley, tomato, and alfalfa leaves. Using knockout mutants, it was shown that biofilm formation impacts on chlorine-disinfection susceptibility after storage (30), and that the biofilm components cellulose and curli impact on transfer and survival (31). On tomato leaflets, a *Salmonella* morphotype that featured extensive biofilm-associated traits attached faster than a morphotype lacking those traits (32). Similarly, *Salmonella* knockout mutants lacking cellulose production featured significantly decreased survival on alfalfa (33). Similarly, the biofilm-forming pathogenic *E. coli* isolates showed decreased susceptibility to sanitizers on leaves of lettuce and baby spinach (34, 35). It is worth noting that bacterial biofilms on produce have been detected and visualized by scanning electron microscopy, in the case of *E. coli* for example on romaine lettuce and spinach leaves (34, 36).

In the present study, we first characterized the *in vitro* biofilm formation capacity of a large collection of 174 environmental *E. coli* strains from irrigation water, fresh produce, soil, or from biofilms scraped from greenhouse sprinklers. Then, we investigated the correlation between *in vitro* biofilm formation and bacterial survival and colonization in the phyllosphere of lamb’s lettuce (*Valerianella locusta*), grown under gnotobiotic conditions. Lamb’s lettuce is one of the most popular salads in Germany (37), and among the most produced salads in Switzerland, accounting for 4,000—5,000 tons per year according to the *Schweizerische Zentralstelle für Gemüsebau und Spezialkulturen* (SZG) (38, 39). We tested the hypothesis that the *in vitro* biofilm formation capacity of *E. coli* isolates is predictive for their persistence on lamb’s lettuce in a gnotobiotic system, serving as a model plant system under controlled laboratory conditions. An extrapolation of the findings from non-pathogenic AR bacteria to pathogens is not intended, since their behaviour can differ significantly.

## Materials and Methods

### Strain origin and characteristics

A total of 174 environmental *E. coli* from different fresh produce obtained at Swiss markets or production fields (n = 35), as well as strains isolated from the production environment of fresh produce (soil (n = 20), irrigation water (n = 117), and greenhouse sprinklers (n = 2)) were tested for biofilm formation capacity *in vitro*. Additionally, full resistance profiles were established by disk diffusion by screening all isolates against 32 antibiotics, covering 14 antibiotic classes (tetracyclines, penicillins, penicillins/β-lactamase inhibitors, 1^st^ to 4^th^ generation cephalosporins, carbapenems, antifolates, quinolones, aminoglycosides, polymyxins, phosphonic antibiotics, and nitrofurans), as described previously (40). Phylogroups of a selected representative number of isolates were determined by PCR and gel electrophoresis according to Clermont *et al.* (41). The multiple antibiotic resistance (MAR) index for each strain was calculated as the ratio between the number of antibiotics to which a strain showed intermediate or full resistance and the total number of antibiotics tested.

For bacterial persistence screening on gnotobiotic plants (*in planta* assays), eleven strains of *E. coli* with different *in vitro* biofilm formation capacity were selected from the 174 *E. coli* strains screened in the *in vitro* biofilm formation assays, while maximizing their diversity regarding antibiotic resistance profiles and isolation source. The eleven strains thereby covered a range of phylogroups, biofilm type, isolation sources, and multiple antibiotic resistance (MAR) indices. Of note, no pathogenic strains or surrogate bacteria were included, as the study focus was on antibiotic resistant bacteria.

### Media and bacterial cultures

Overnight cultures were grown in lysogeny broth (LB; Sigma-Aldrich, St. Louis, USA) at 37°C if not otherwise indicated. For crystal violet (CV) assays, LB-NaCl(5) (10 g/l peptone, 5 g/l yeast extract, 5 g/l NaCl, pH 7.0), LB-NaCl(0) (10 g/l peptone, 5 g/l yeast extract, pH 7.0), and AB minimal medium (1µg/mL thiamine, 25 µg/mL uridine, and 10 µg/mL proline) with 0.5% w/v casamino acids as carbon source (ABTCAA) (42, 43) were used.

For selective culturing of *E. coli*, CHROMagar *E. coli* (CrA; CHROMagar^TM^; Paris, France) was used. To select for antibiotic-resistant strains, CrA supplemented with either ampicillin (AM, 100 mg/L), kanamycin (K, 16 mg/L), ciprofloxacin (CIP, 1 mg/L), or ceftazidime (CAZ, 8 mg/L) was used. Unless otherwise indicated, CrA plates were incubated for 24 h at 37°C.

### Crystal violet assays

To quantify biofilm formation *in vitro*, a 96-well plate-based system was used as described by Marti *et al.* (44), which relies on the colorimetric quantification of crystal violet. Flat bottom, 96-well polystyrene microtiter plates (CytoOne^®^; USA Scientific, Ocala, USA) were used for all assays. Four technical replicates were recorded per strain and the experiments were repeated three times independently. Finally, two reference strains, *E. coli* ATCC 25922 and *E. coli* FAM 21843, a weak and strong biofilm formation control, respectively (45), as well as non-inoculated medium (negative control) were measured on each 96-well plate in quadruplicate.

To perform the CV assay, strains were grown overnight in LB-NaCl(5) at 37°C. Cultures were then diluted 1:100 in either LB-NaCl(0) or ABTCAA, representing a nutrient-rich or - poor medium, respectively. Notably, ABTCAA is chosen as a medium with minimal nutrient availability (as opposed to nutrient-rich LB), to assess the effect of nutrient availability on biofilm formation. Per treatment, four wells were filled with 150 μL of diluted culture; the microtiter plate was placed in a polypropylene (PP) plastic incubation chamber, and incubated for 48 h at 28°C. After incubation, the medium was removed gently and attached cells were washed three times with 200 μL/well of sterile physiological saline (0.9% w/v NaCl). Then, 200 μL 0.1% v/v CV solution (Sigma-Aldrich, St. Louis, USA) per well were added, plates were covered with a lid, and incubated at room temperature for 20 min to stain biofilms. The CV stain was then discarded and the stained biofilms were washed three times with 200 μL/well of sterile distilled water (Milli-Q^®^) to remove excess stain. Finally, the biofilm-absorbed stain was dissolved in 200 μL/well of 96% v/v ethanol (Sigma-Aldrich), and optical density was recorded at 600 nm (OD_600_) using a Tecan Infinite M200 microplate reader (Tecan Group AG, Männedorf, Switzerland). Of note, when the measured OD_600_ exceeded 1.0, 10- or 20-fold dilutions were prepared in 96% v/v ethanol and re-measured.

To select *E. coli* strains for the gnotobiotic plant system, strains were assigned to biofilm formation categories according to Stepanović *et al.* (46). Briefly, a cut-off value (OD_c_) was defined as three standard deviations (SD) above the mean OD_600_ of the negative control (non-inoculated medium). The following categories were then defined as: none/weak biofilm formation (OD_strain_ ≤ 2×OD_c_), moderate biofilm formation (2×OD_c_ < OD_strain_ ≤ 4×OD_c_), and strong biofilm formation (4×OD_c_ < OD_strain_ ≤ 10×OD_c_). Since the tested *E. coli* collection encompassed some extreme biofilm formers, we defined an additional category of extreme biofilm producers (10×OD_c_ < OD_strain_).

### Gnotobiotic plant system

#### Seed sterilization

Lamb’s lettuce (*Valerianella locusta*) was used to establish an *in vitro* gnotobiotic plant growth system. The rational for using a gnotobiotic system –which differs significantly from the field or market non-sterile produce– was to achieve a highly controlled and reproducible environment over a large experimental setup, as well as to reduce the variability between batches for improved comparability. Gnotobiotic systems allow to minimize artifacts and introduce specific isolates onto plants to study their behaviour without the unwanted effect of other agents, as argued in many studies (47–50). Thereby, the plant microbiome otherwise present does not have to be considered. To obtain sterile plantlets, *V. locusta* seeds (var. ‘dark green full rosette’, Select, Wyss Samen und Pflanzen AG, Zuchwil, Switzerland) were sterilized as follows: sterilization solution was prepared from sodium hypochlorite (NaOCl) with 11–14% v/v available chlorine (Alfa Aesar, Haverhill, Massachusetts, USA) by diluting it 1:3 with sterile distilled water to obtain a 3% v/v bleach solution. Two hundred milligrams of seeds were then washed with 3.2 mL of sterile distilled water and harvested by centrifugation (1 min, 1,500 × *g*). After discarding the liquid, seeds were washed with 3.2 mL of 70% v/v ethanol for 1 min, centrifuged again (1 min, 1,500 × *g*), and the liquid was again discarded. Then, 3.2 mL of sterilization solution were added and the seeds were vortexed for 2 min. After centrifugation and liquid removal, the seeds were washed three times with sterile distilled water by consecutive vortexing, centrifugation, and removal of the liquid. The seeds were then transferred to a glass petri dish and allowed to dry under laminar flow for approximately 30 min. Finally, the petri dish was sealed with parafilm and incubated at 10°C for 7 days in the dark for seed stratification.

#### Seed sprouting and planting

The sterile seeds were allowed to sprout in fluted filter paper designed to that purpose (0.22 mm thickness, 2,000 × 110 mm, #146036; Munktell Filter AB, Falun, Sweden): The fluted filter and the corresponding wrapping filter (#123420) were placed inside a sterile glass dish (ø 22 cm), and three to four seeds were placed into each filter fold. The fluted filter was then covered with the wrapping filter, watered with 50 mL of sterile tap water, and the covered/parafilm-sealed glass dish was placed into the plant incubation chamber (12 h at 21°C and 12 h at 17°C). Note that the seeds were kept in the dark for 7 days before switching to a day-night-cycle with 12 h of light for another 7 days.

After the 2-week incubation in the filter, seedlings were transferred aseptically into UV-sterilized, transparent LDPE plastic boxes (diameter: 8 cm (bottom) to 9 cm (top), height: 6 cm) containing 100 mL of half strength Murashige-Skoog (MS) agar (2.2 g/L MS salts from Duchefa, 10 g/L sucrose, and 5.5 g/L plant agar from Carl Roth, Karlsruhe, Germany; pH 5.8). Before seedling transfer, four crosses were cut deeply into the agar with a sterile scalpel to facilitate planting. The sealed bowls were then transferred to the plant chamber and incubated for additional 21 days.

#### Inoculation and harvest

Single *E. coli* colonies were picked from CrA and streaked over a whole LB agar plate to obtain a bacterial lawn after incubation for 24 h at 37°C. Bacterial cells were harvested with a 10 μL loop and suspended in 3 mL of phosphate-buffered saline (PBS, 8 g NaCl, 0.2 g KCl, 1.15 g Na_2_HPO_4_, and 0.2 g KH_2_PO_4_ in 1 L distilled water; pH 7.3). After thorough resuspension by vortexing, bacterial suspensions were centrifuged (4,500 × *g*, 5 min) and the supernatant discarded. The pellet was resuspended in 10 mL of PBS, OD_600_ was measured, and appropriate dilutions were prepared to obtain an OD_600_ of 0.05 ± 0.005, which was used for plant inoculation. The four plants per box were each inoculated with 10 μL of OD-adjusted culture of one strain by distributing small droplets on the first two rosette leaf pairs, and plants were returned into the incubation chamber. Bacterial counts at inoculation (0 days post inoculation; 0 dpi) were determined by plating appropriate dilutions of the OD-adjusted cultures on LB as well as CrA agar, to detect potential contamination by comparing the non-selective and selective agar counts.

To determine bacterial survival in the phyllosphere, plants were harvested at 3, 7, 14, and 21 dpi. Six plants per strain and time point were harvested from six different boxes. Additionally, one negative control plant (inoculated with sterile PBS) was harvested at each sampling time. The whole time series experiment was repeated three times independently. For harvesting, plants were picked with sterile tweezers and the roots were cut off with a scalpel before transferring the plant into a 5 mL Eppendorf tube and determining the plant fresh weight. Twenty glass beads were then added into each tube along with 1 mL of PBS. Plants were then bead-beaten horizontally on a vortex for 15 min to detach bacteria from the leaves. Each leaf macerate and its dilutions were then plated on LB agar as well as CrA to detect contaminating bacteria on the plants by comparing selective- and non-selective medium counts. Plants showing significant deviation of cell counts between LB agar and CrA (above one order of magnitude) were excluded from analysis. Of note, plating of the negative plants revealed an acceptable efficiency of sanitation with only 6 out of 76 negative plants displaying growth of single colonies on LB, but no growth on CrA. The lower limit of detection was 5 CFU/mL of leaf macerate.

### Fluorescence microscopy

Fluorescence microscopy was conducted on an Axioplan microscope (Carl Zeiss AG, Feldbach, Switzerland) at 20× or 40× magnification (EC Plan-Neofluar 20×/0.50 Ph 2 M27 or EC Plan-Neofluar 40×/0.75 Ph 2 M27 objectives, respectively), equipped with a high-pressure mercury arc lamp (HBO 50; Carl Zeiss AG), the Zeiss filter sets 38HE (BP 470/40-FT 495-BP 525/50) and 43HE (BP 550/25-FT 570-BP 605/70), an AxioCam MRm3 and the software Zeiss ZEN 2.6 (Blue edition). To visualize bacterial cells on the leaf surfaces, the dye LIVE/DEAD BacLight Kit L7007 (Molecular Probes Inc., Eugene, USA) was used following the manufacturers recommendations. After 7, 14, and 21 dpi two single leaves were harvested per strain and time point, and 20 μL of diluted dye were pipetted onto each leaf before covering it with a cover slip (24 × 50 mm; Menzel-Gläser, Braunschweig, Germany) and fixing the slide with adhesive tape. After 15 min incubation in the dark at room temperature, the leaves were ready for microscopy. A minimum of four representative field of views per leaf were recorded. The control strain *E. coli* C13, a moderate biofilm former originally isolated from reservoir water, as well as two leaves from a plantlet inoculated with PBS (negative control) were imaged in every round. Microscopic pictures were processed using ImageJ2/FIJI v2.3.0/1.53f. Z-stacks were first combined using the maximum intensity Z-projection function, in which the resulting images were corrected by applying a rolling ball background subtraction algorithm (50 px) and contrast enhancement (0.1%). Composite images were generated and single channels were assigned a pseudo-colour for visualization (Yellow: live cells; Magenta: dead cells). To determine representative images, we manually screened our collection of 365 pictures and selected an appropriate number of images, representing the different bacterial growth behaviors.

### Quantitative PCR

Biofilm formation can be expected to affect the accuracy of bacterial plate counts, as aggregated bacterial cells, rather than single bacterial cells form individual colonies, which may result in underestimation of CFU. We therefore used quantitative PCR (qPCR) additionally to plate counting, and investigated the correlation between CFU counts and *yccT* copy numbers for all strains used on gnotobiotic plants.

Quantification of viable *E. coli* was done by viability qPCR after propidium monoazide (PMA) treatment, to avoid the quantification of cells with compromised cell membranes, which are susceptible to DNA modification by the stain and are considered as dead cells (51). To that end, 500 µL of leaf macerate were treated with 25 µM of PMAxx (Biotum, Fremont, USA) and incubated with shaking at 400 rpm, at room temperature in the dark for 10 min. Then, samples were exposed for 15 min to a 60 W LED light to promote the covalent binding of PMAxx to the dsDNA of damaged cells, resulting in permanent DNA modification that prevents PCR amplification. Afterwards, PMA-treated cells were concentrated by centrifugation at 6,000 × *g* for 10 min. The resulting cell pellet was resuspended in 100 µL distilled water and DNA was extracted. Briefly, the cell suspension was heated to 95°C for 10 min, then cooled down at −20°C for 10 min and finally, the DNA was recovered from the supernatant after centrifuging at 10,000 × *g* for 5 min. Viable *E. coli* were quantified by qPCR targeting the *yccT* gene, which is specific to *E. coli* (52). The assays were performed on a Rotor-Gene thermocycler (Qiagen, Hombrechtikon, Switzerland) in a 25 µL reaction including 1x SureFast® Master Probe PLUS (Congen, Berlin, Germany), 0.2 µM of forward (5’-GCATCGTGACCACCTTGA-3’) and reverse (5’- CAGCGTGGTGGCAAAA-3’) primers, 0.1 µM of TaqMan probe (5’- TGCATTATGTTTGCCGGTATCCG-3’), and 2 µL of DNA template. The primers and probes used in this study were previously designed (52). Cycling conditions included a first step at 95°C for 10 s and 40 cycles of denaturation at 95°C for 10 s followed by annealing/extension at 56°C for 15 s. To quantify the number of bacterial cells in the samples, a standard curve consisting of 5-log DNA dilution series from a known concentration of control strain *E. coli* C13 was included in duplicate for every qPCR assay.

### Data analysis and statistics

Data processing, statistical analysis, and graphical representations were performed in R (53). Data organisation and visualisation were performed using the *tidyverse* package (54).

*In vitro* biofilm formation data was analysed using the log_2_-transformed OD_600_ values from the biofilm assays. To test for differences in OD_600_ values of strains grown in two different media, a paired t-test was performed with the *t.test()* function of the *stats* package. Linear mixed-effect (LME) regression analyses were done with the *lme()* function of the *nmle* package (55). Factors such as type of medium (ABTCAA, LB-NaCl(0)), source of isolation (fresh produce, soil, water), or phylogroup were used as explanatory variables in the regression analyses, and strain identity or medium were used as grouping factor (random effects), depending on the model. The effect of each variable was evaluated with an analysis of variance on the fitted model using the *anova()* function of the *stats* package. Outlier detection for *in planta* CFU counts and qPCR data was conducted by estimating Cook’s distances for each data point. A cutoff of four times the mean Cook’s distance (*cooks.distance()* from the *stats* package) was applied to detect and filter out outliers from downstream analyses (46 out of 1157 measurements).

Pearson’s correlations were calculated using the *cor()* function from the *stats* package, to determine the correlation between *in vitro* OD measurements in rich and minimal media, as well as the correlation between CFU and qPCR measurements of bacterial population densities on plants.

For *in planta* bacterial persistence analysis, the log_10_-transformed data of the *yccT* gene abundance was used. To account for differences in variances between independent experiments and sampling days, a generalised least squares (GLS) model was used to explain changes in *yccT* gene copy numbers by different explanatory variables (e.g., time, biofilm type, or strain identity), using the *gls()* function of the *nlme* package. Different variance structures were tested and model selection was performed according to the Akaike information criterion (AIC). The effect of factors and interactions were determined through Wald tests performed with the *anova.gls()* function of *nmle*. For every model and when appropriate, pairwise comparisons were made by comparing least-squares means of each factor with the package *emmeans* (56).

To better describe the sample distribution of each *E. coli* strain grown *in planta* at different time points, population densities (log_10_-transformed data of the *yccT* gene abundance) were further analysed as mixtures of two Gaussian distributions using the *mixtools* package (57). Posterior distributions were plotted with *ggplot*.

## Results

### *E. coli* biofilm formation *in vitro*

#### Medium type affects biofilm formation

We observed that biofilm formation depended on the media used (LB-NaCl(0): rich; ABTCAA: poor), although the effect was small (Fig 1A). In LB-NaCl(0), OD_600_ values were 1.4 times higher than in ABTCAA (paired t-test, *t*(173) = 5.24, *p* < 0.05). Consequently, the biofilm type assignment changed for a number of strains. From the 174 strains tested, 54 strains showed increased OD_600_ values in LB-NaCl(0) compared to ABTCAA (green lines, Fig 1A), resulting in 4 strains changing from none/weak to moderate or strong, 3 from moderate to strong or extreme, and 47 from strong to extreme biofilm formation. Of the main isolation sources (fresh produce, soil, and water), this increase was most frequent in water isolates (34.2%; Tab S1).

**Figure 1.**
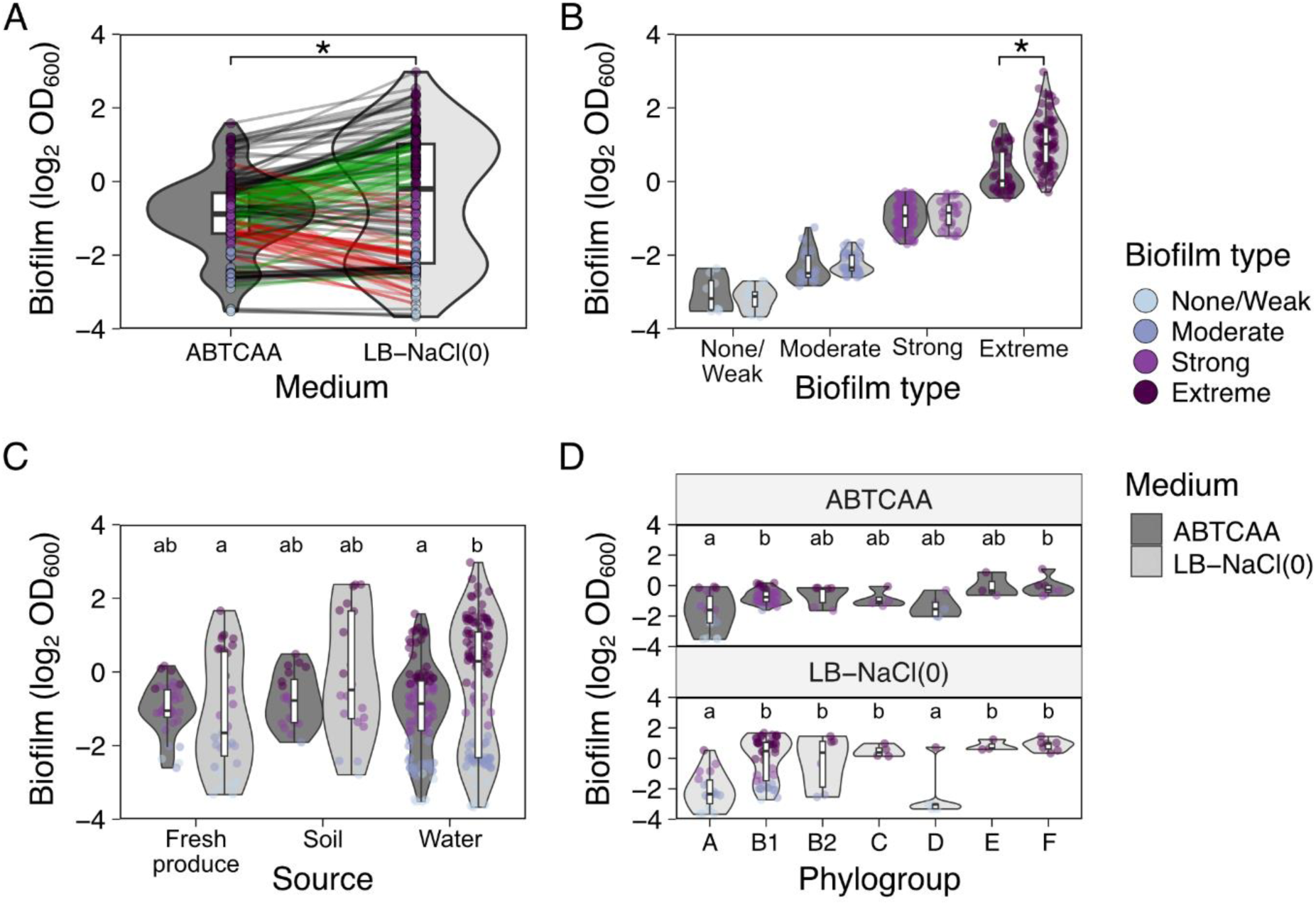
*In vitro* biofilm formation capacity as determined by crystal violet (CV) staining and quantification of CV stain by OD measurement at 600 nm. (A) Biofilm formation capacity by medium. Changes in biofilm type of individual strains depending on the medium are highlighted with coloured lines (green: change from ABTCAA [AB minimal medium with casamino acids] to LB-NaCl(0) [nutrient rich medium without NaCl] to a stronger biofilm type, red: change from ABTCAA to LB-NaCl(0) to a weaker biofilm type, black: no change in biofilm type between media). (B) Biofilm formation capacity by biofilm type (none/weak, moderate, strong, and extreme) in the two media. In A and B: asterisk indicate statistically significant differences between with 𝛼 = 0.05. (C) Biofilm formation capacity by source of isolation and by medium. Letters indicate statistically significant differences between groups (source and media) with 𝛼 = 0.05. (D) Biofilm formation capacity by a representative number of phylotyped isolates relative to their phylogroup and growth medium. Letters indicate statistically significant differences between phylogroups within a medium, with 𝛼 = 0.05, from linear mixed-effect regression models. In all cases, violin plots are depicted to show the distribution of the data and boxplots indicating the minimum, the interquartile range, and maximum value. A strain-specific representation of the data can be found on [Zenodo](https://zenodo.org/records/13838842).

By contrast, 41 strains decreased their OD_600_ values in LB-NaCl(0) compared to ABTCAA (red lines, Fig 1A), resulting in 5 strains changing from moderate to none/weak, 29 from strong to moderate or none/weak, and 7 strains from extreme to strong or moderate. Of the main isolation sources, this decrease was most frequent in fresh produce isolates (48.6%; Tab S1).

The OD_600_ values used to categorise strains in their *in vitro* biofilm formation capacity were similar between the two media, except for the extreme biofilm type, for which significantly higher OD_600_ values were measured in LB-NaCl(0) compared to ABTCAA (Fig 1B; *F*_3,166_ = 20.79, *p* < 0.05). Overall, a strong correlation was found between biofilm formation capacities in ABTCAA and LB-NaCl(0) (Fig 2; Pearson’s *r* = 0.71, *t*(172) = 13.3, *p* < 0.05).

**Figure 2.**
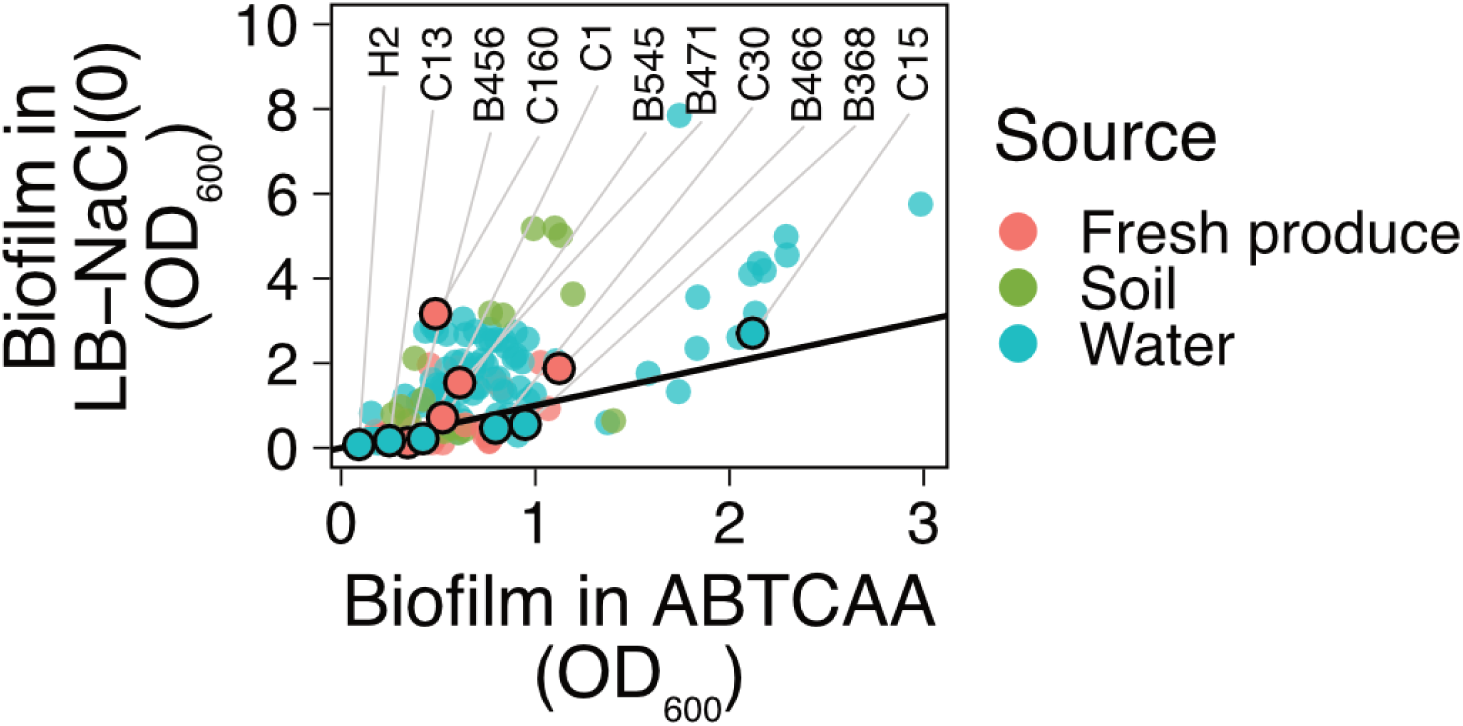
Correlation between bacterial biofilm formation capacity as determined by crystal violet (CV) staining and quantification of CV stain by OD measurement at 600 nm, in ABTCAA [AB minimal medium with casamino acids] and LB-NaCl(0) [nutrient rich medium without NaCl]. Pearson’s correlation, *r* = 0.71. Highlighted strains designated with letter and number (e.g., H2) were selected for bacterial persistence screening on gnotobiotic *V. locusta* plantlets. As a reference, the identity function is shown as a continuous line: strains below or above the line showed stronger *in vitro* biofilm formation in ABTCAA or LB-NaCl(0), respectively.

#### Mean *E. coli in vitro* biofilm formation weakly correlates with isolation sources and phylogroups

Considering variables other than growth medium, *in vitro* biofilm formation differed slightly between strains from different isolation sources (fresh produce, soil, or water). We excluded greenhouse sprinkler isolates from this analysis because of the low sample number (n = 2). Although differences were small (1.2—2.1 fold), strains isolated from fresh produce showed lower mean OD_600_ values compared to strains isolated from soil or water. These differences were only observed for biofilm formation capacity determined in LB-NaCl(0) (Fig 1C; *F*_2,169_ = 6.69, *p* = 0.0016). Notably, LB-NaCl(0) seemed to encourage a bimodal distribution, as is especially evident in water isolates (Fig 1C).

When evaluating the relationship between phylogroups with *in vitro* biofilm formation capacity, we observed that some phylogroups significantly differed in their ability to form biofilms depending on the growth medium (Fig 1D; *F*_6,79_ = 3.92, *p* = 0.0018). Particularly, phylogroup A strains were consistently different from phylogroup B1 and F, regardless of the media.

### *In planta* persistence of AR *E. coli* of different biofilm types

#### Strain characteristics

Combined, the selected isolates represent the most common *E. coli* phylogroups (A, B1, D, and F) and covered the whole range from none/weak to extreme biofilm formers. They originated from a variety of water sources including surface, drainage, reservoir, and sprinkler irrigation water, as well as from different fresh produce (Fig 3). Finally, they were all multidrug-resistant according to the definition by Magiorakos *et al.* (58), including an ESBL-producing strain H2 (Fig 3). The highest observed MAR index, 0.66, was exhibited by strain H2, while the lowest MAR index, 0.09, was exhibited by strain C15 (Fig 3). However, there was no strong correlation between *in vitro* biofilm formation and MAR in ABTCAA (Fig S2; *r* = −0.57, *t*(9) = −2.09, *p* = 0.066) or LB-NaCl(0) (*r* = −0.43, *t*(9) = −1.43, *p* = 0.19).

**Figure 3.**
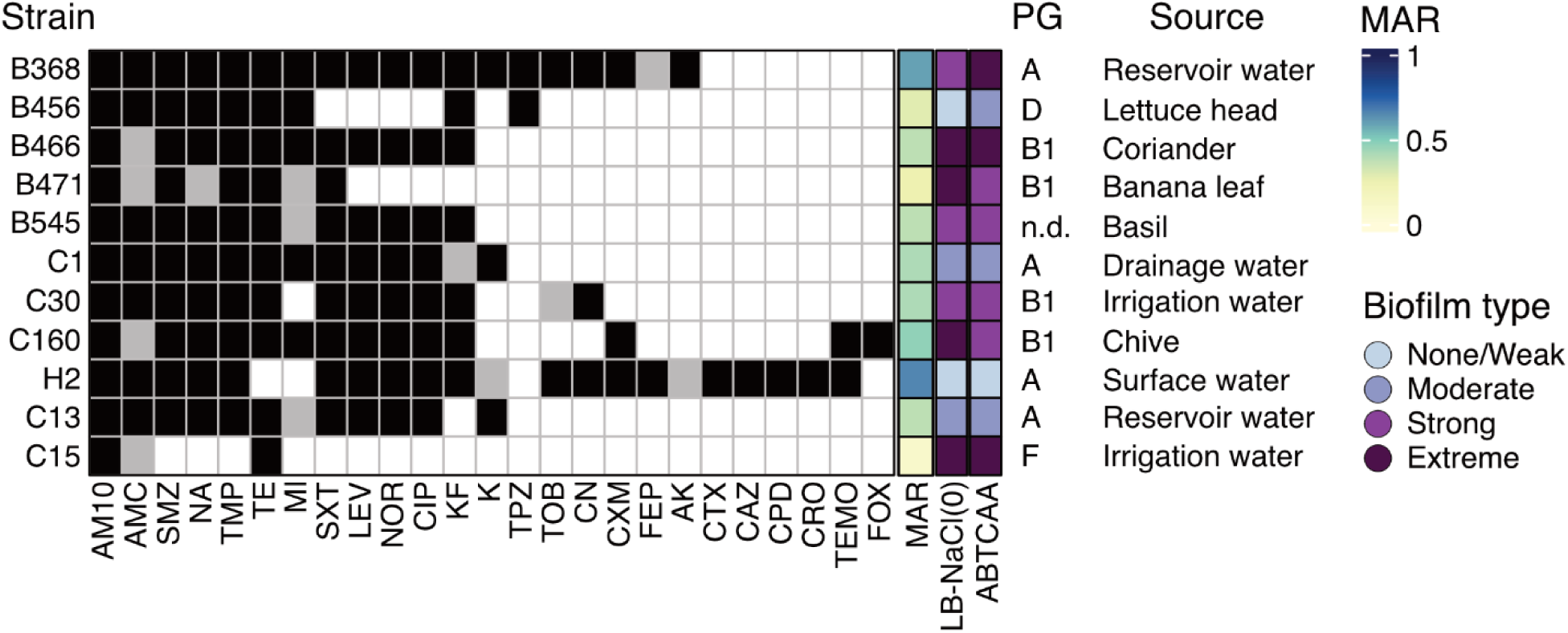
Characteristics of bacterial strains selected for *in planta* assays. Antibiotic resistance profile, multiple antibiotic resistance (MAR) index, biofilm type in ABTCAA [AB minimal medium with casamino acids] and LB-NACl(0) [nutrient rich medium without NaCl], phylogroup (PG), and isolation source are indicated for each strain. Black: resistant; light grey: intermediate; white: sensitive. AM10: ampicillin 10 μg; FEP: cefepime; FOX: cefoxitin; CPD: cefpodoxime; AMC: amoxicillin-clavulanic acid; CRO: ceftriaxone; TPZ: piperacillin-tazobactam; CXM: cefuroxime; TOB: tobramycin; CN: gentamicin; NA: nalidixic acid; NOR: norfloxacin; SXT: trimethoprim-sulfamethoxazole; MI: minocycline; K: kanamycin; AK: amikacin; CIP: ciprofloxacin; LEV: levofloxacin; TMP: trimethoprim; SMZ: sulfonamide; TE: tetracycline; TEMO: temocillin; KF: cefalotin; CAZ: ceftazidime; CTX: cefotaxime. All strains were sensitive towards tigecycline, ertapenem, meropenem, imipenem, nitrofurantoin, fosfomycin, and colistin; *n.d.*: not determined. MAR index was calculated based on the screening of 32 antibiotics.

#### High correlation between CFU counts and *yccT* copy numbers

Biofilm formation can be expected to affect the accuracy of bacterial plate counts, as aggregated bacterial cells, rather than single bacterial cells form individual colonies, which may result in underestimation of CFU. We found a strong correlation between the two quantification methods, regardless of *in vitro* biofilm formation capacity or strain (Pearson’s *r* = 0.89, *t*(1109) = 63.9, *p* < 0.05; Fig 4A). Among biofilm-forming strains, moderate and strong biofilm producers showed a higher correlation between *yccT* copy numbers and CFU per gFW (Pearson’s *r* = 0.90 and 0.89, respectively; Fig 4B) compared to none/weak and extreme biofilm formers (Pearson’s *r* = 0.85 and 0.86; Fig 4B). Regression analysis revealed that for none/weak and extreme biofilm-forming strains, the slope coefficients differed significantly from 1 (*β*_none/weak_ = 0.92, *Z* = 2.22, *p* = 0.026; *β*_extreme_ = 0.89, *Z* = 2.85, *p* = 0.0044, respectively; Fig 4B), indicating a deviation between CFU counts and qPCR data.

**Figure 4.**
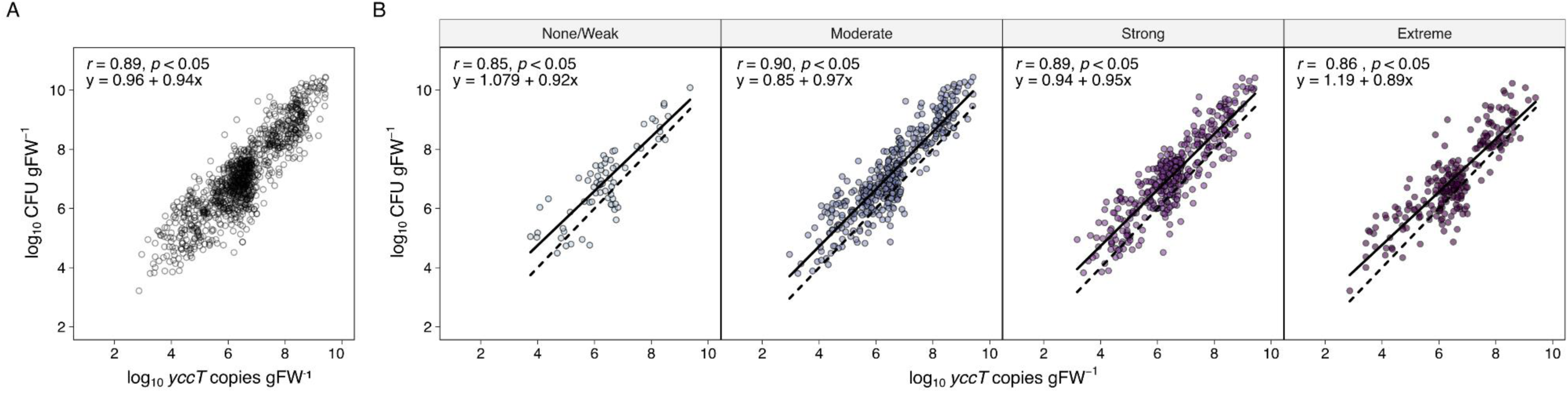
Correlation between bacterial culture data (CFU per g fresh weight (gFW)) and qPCR data (*yccT* copies per gFW) for each AR *E. coli* strain tested on gnotobiotic *V. locusta*, (A) all the data combined or (B) grouped by biofilm type (none/weak, moderate, strong, extreme) determined in ABTCAA medium [AB minimal medium with casamino acids], at every sampling time point and from all conducted experiments (log_10_ scale). Pearson’s correlation (*r*), p-values (*p*), and linear equations are displayed. In B, continuous line represents the linear relationship between observed values, while dashed line represents a reference line of perfect equivalency (y = x).

The bead beating method applied for detaching bacterial cells from the leaves prior to plating therefore seems efficient in disrupting cell clumps. Following the good correlation between the two quantification methods over all investigated strains (Fig 4), data analysis was based solely on qPCR data of PMA treated samples as it is expected to be more accurate than plate counting.

#### *In vitro* biofilm formation does not affect *in planta* bacterial population densities

We tested whether the differences in *yccT* gene copy numbers, i.e., population density, between strains *in planta* were explained by the biofilm formation capacity *in vitro* and the time since plant inoculation. As a reference, we used the strain’s *in vitro* biofilm capacity in ABTCAA medium providing poor resource conditions (44), as the phyllosphere environment is considered to be oligotrophic (19). We observed that differences depended on the interaction between *in vitro* biofilm capacity and time (Fig 5A; *F*_12,1091_ = 3.55, *p* < 0.05), although the effect size was negligible (Cohen’s *f^2^* = 0.026). Similarly, an interaction effect between independent strains and time was observed (Fig 5B; *F*_40,1056_ = 2.04, *p* = 0.0002), with a small effect size (Cohen’s *f^2^* = 0.050). The only differences between strains were identified at 21 dpi (Fig 4B), with the *in vitro* extreme biofilm-forming strain C15 showing a larger mean population size than B466 (extreme), C160 and C30 (moderate), and H2 (none/weak).

**Figure 5.**
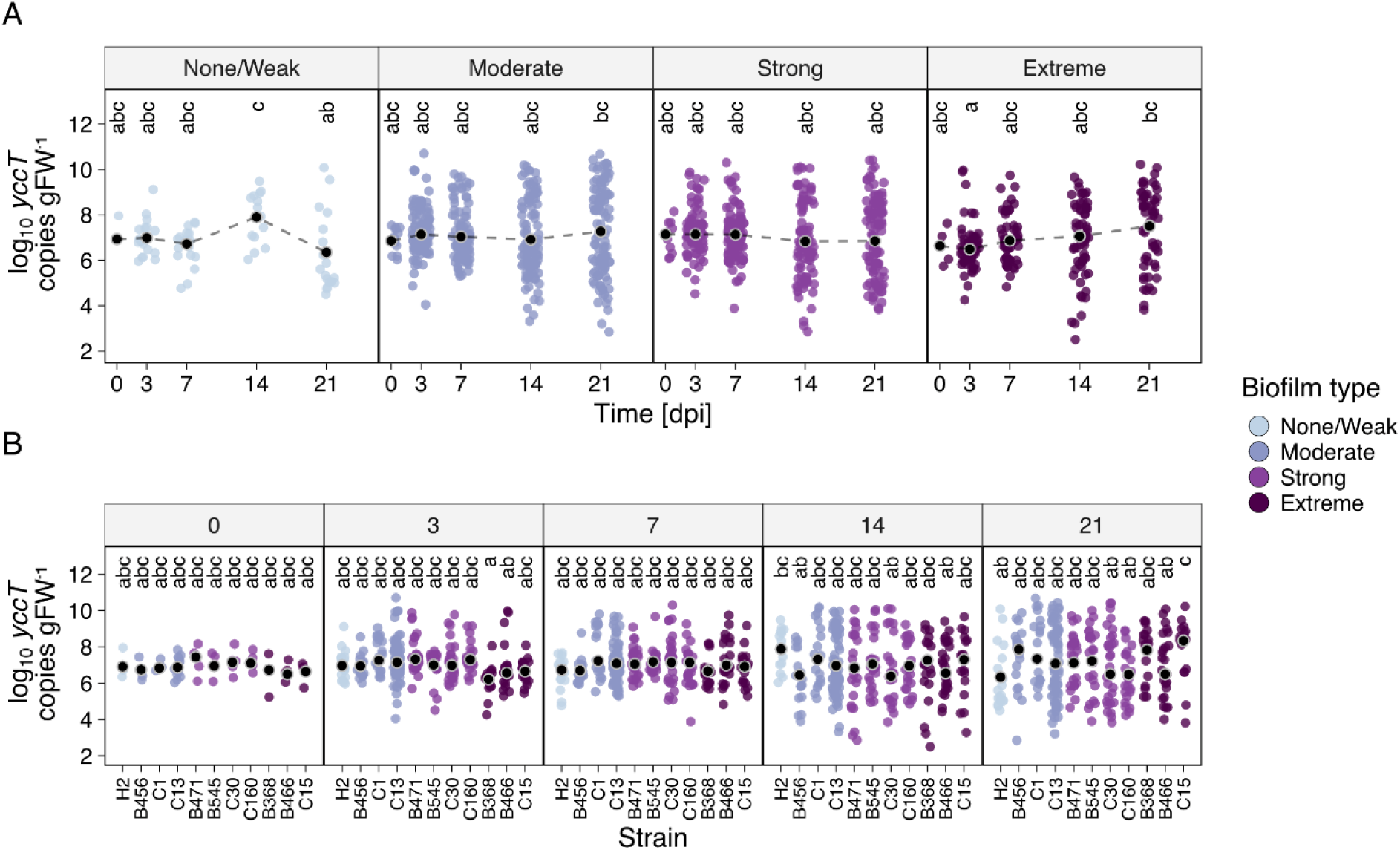
Bacterial persistence over time on leaves of gnotobiotic *V. locusta*. Each point depicts the AR *E. coli* abundance as copies of the *yccT* gene on each replicate plant at a given sampling point (n = 6 plants per strain and time point; N = 3 independent experiments). The outlined black point indicates the estimated marginal mean for a group from a generalised least square model. Strains were grouped according to biofilm type determined in ABTCAA medium [AB minimal medium with casamino acids] (A), or shown as individual strains (B). Groups with different letters indicate significant differences from adjusted multiple pairwise comparisons (*p* < 0.05). gFW, gram fresh weight; dpi, days post inoculation.

In addition, we tested whether MAR could have a predictive effect on population density. However, while we observed an interaction between MAR and type of biofilm-forming strain (Fig S4; *F*_2,1143_ = 6,59, *p* = 0.0014), the effect was low as it explained only 1.12% of the total variance.

#### Time-dependent distinction of two subpopulations *in planta*

While *in vitro* biofilm formation capacity did not affect a strain’s capacity to survive on the gnotobiotic plants, we noticed that for moderate, strong, and extreme biofilm-forming strains, the dispersion of population densities on the plants increased over time, clearly distinguishing two subpopulations at 14 and 21 dpi (Fig 5A and 5B). Since these subpopulations were characterised by a seemingly bimodal distribution, an expectation-maximisation algorithm was used to fit a mixture of Gaussian distributions with two components, resulting in an improved fit of the *yccT* gene copy data than when considering the distribution of a population as a whole (Fig 6). Especially at 14 and 21 dpi, two subpopulations could be detected for each biofilm-forming strain type. In other words, each of the strains of the above-mentioned *in vitro* biofilm types was able to grow and establish well on certain plants, while on other plants its population density would decrease over time. In confirmation of this observation, the coefficient of variation of strains increased over time for each biofilm group (Fig S3A) as well as for each individual strain (Fig S3B), suggesting an increase in the variability of bacterial abundance on *V. locusta*. Additionally, we observed this same pattern across independent experiments, each of which included strains of different biofilm types (Fig S3C).

**Figure 6.**
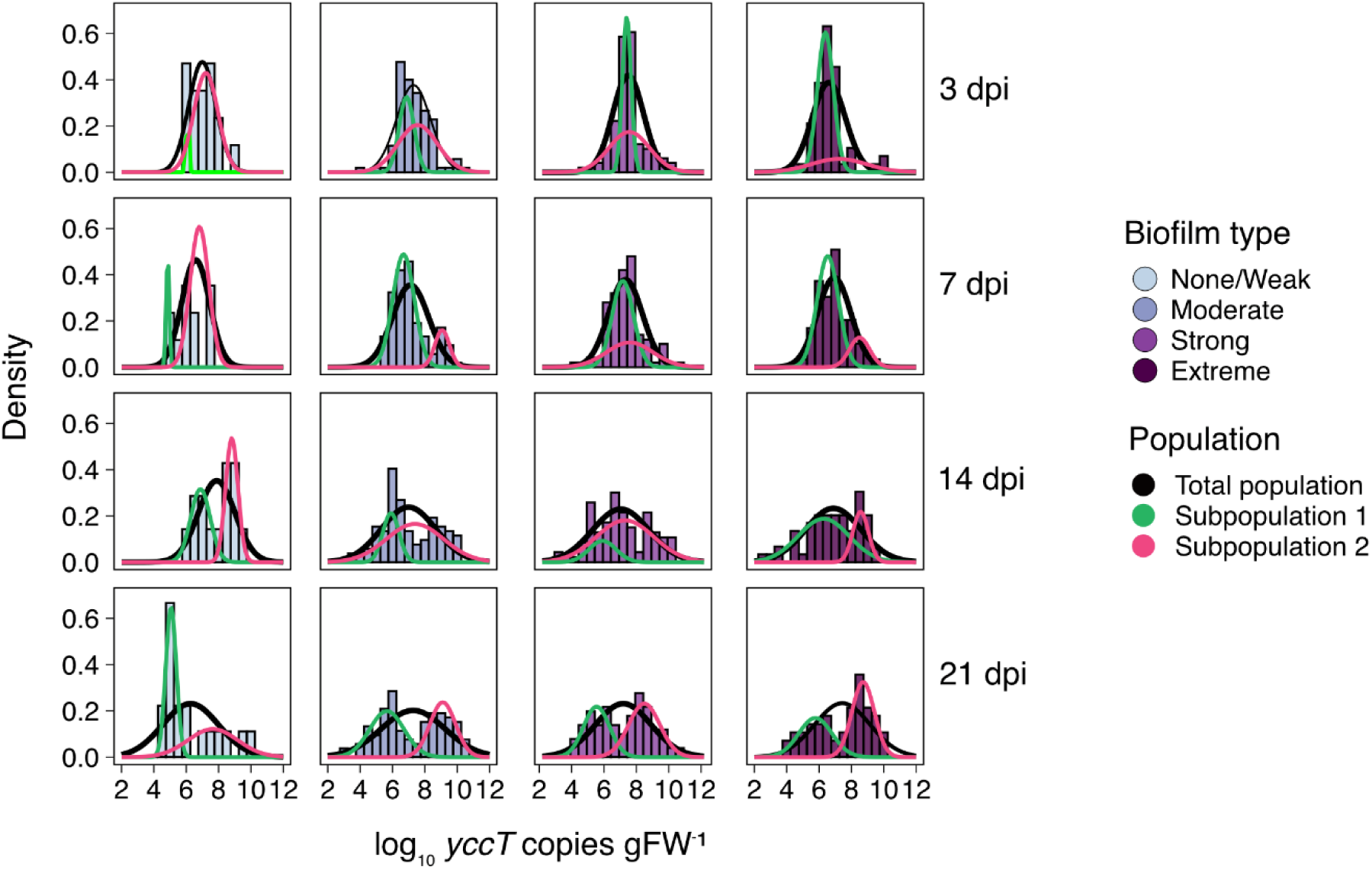
Distribution of biofilm-forming *E. coli* populations on gnotobiotic *V. locusta*. Histograms and probability density function of the abundance of *E. coli* population as *yccT* copies per gram of fresh weight (gFW^-1^) over time (dpi, days post inoculation) for strains grouped by biofilm type determined in ABTCAA medium [AB minimal medium with casamino acids]: none/weak, moderate, strong, and extreme. Total population distribution is depicted as a black line, while subpopulations derived from mixture models are shown as green and magenta lines.

#### *Escherichia coli* forms long aggregates on *V. locusta* leaves

To inspect colonization of gnotobiotic *V. locusta* leaves by *E. coli* strains of different *in vitro* biofilm types visually, a time series of fluorescence microscopy pictures was recorded per strain. One representative picture was selected per time point for one strain of each biofilm type (Fig 7). Overall, microscopy confirmed the observations from the bacterial culture and *yccT* gene copy data, that is, no striking differences in colonization efficiency and persistence on the gnotobiotic plant leaves were observed between strains of different biofilm types, although different colonization types were observed. Strains of all biofilm types were able to form long aggregates that follow epidermal cell grooves (e.g., strain B466: “extreme” in Fig 7), while other strains populated the leaves preferably as single cells or in small colony-like structures (e.g., strain H2: “none/weak” in Fig 7).

**Figure 7.**
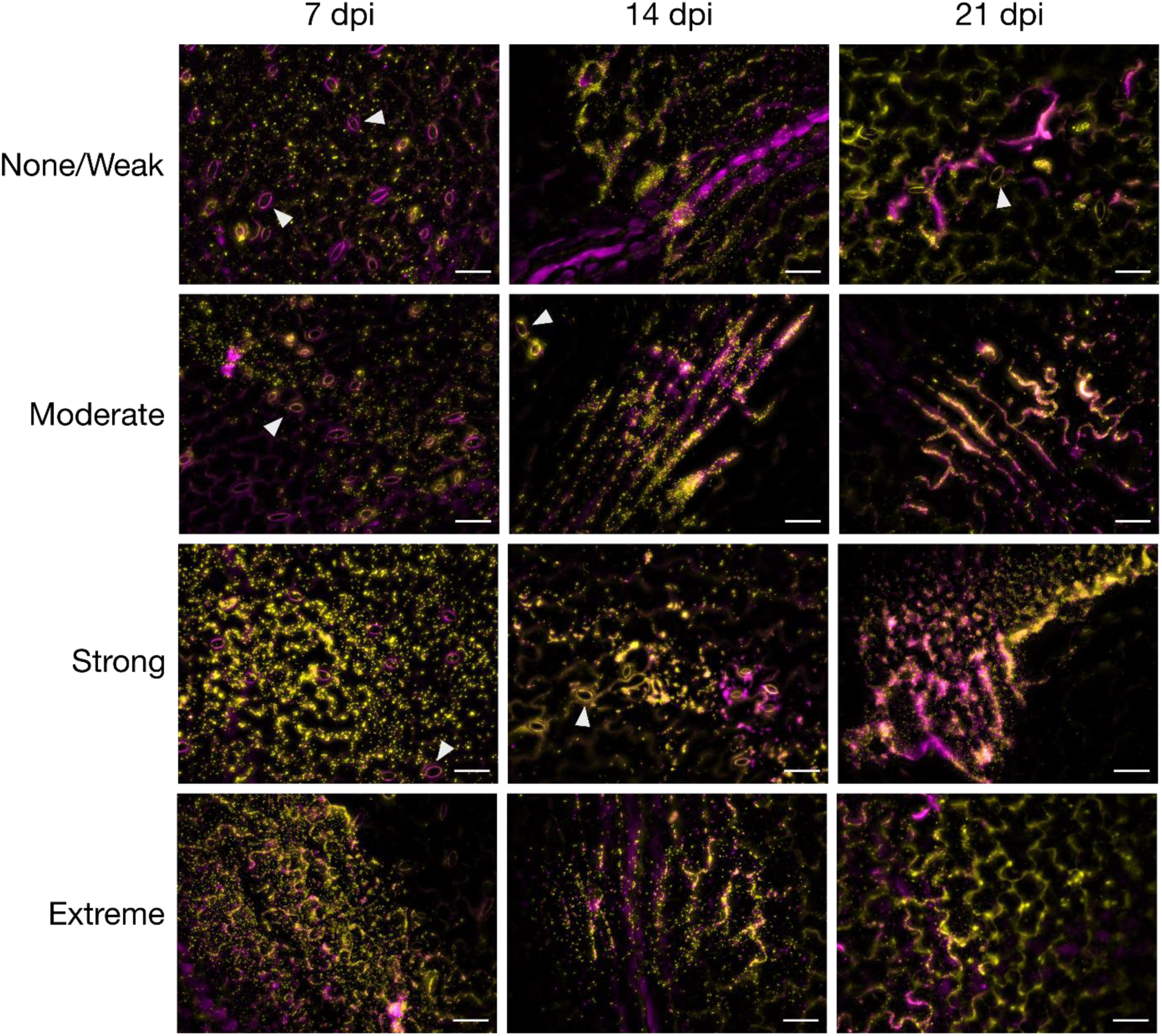
Live/dead stain micrographs of representative AR *E. coli* of different biofilm types determined in ABTCAA medium [AB minimal medium with casamino acids], colonizing the phyllosphere of gnotobiotic *V. locusta*. Images were taken at 7, 14, and 21 days post inoculation (dpi) of the plantlets. Images were the result of overlaying the live/dead fluorescence channels: yellow correspond to viable cells; magenta correspond to not viable cells and plant autofluorescence. Biofilm types and corresponding representative strains: none/weak: strain H2; moderate: strain C13; strong: strain C30; extreme: strain B466. Scale bar = 50 µm. Triangles indicate plant leaf stomata.

## Discussion

Biofilm formation is a common strategy of bacterial cells to persist in the environment, and it has often been considered a key characteristic of bacterial strains along with their resistance and virulence traits. However, it has been recognized that findings from *in vitro* biofilm models often differ significantly from what is observed *in vivo* (59). A validation of *in vitro* results using *in vivo* systems therefore is key to draw conclusions for real-world systems. Nevertheless, such comparative studies are relatively rare (60), as many studies use *in vitro* models only.

In the present work, we observed that the type of growth medium affected *in vitro* biofilm formation capacity for over half of the examined environmental *E. coli* isolates. The remaining strains did not change their biofilm type when grown in minimal instead of rich medium. It has been observed that biofilm formation is inhibited by catabolite repression, possibly an indicator of a nutrient-rich environment, where growing in biofilms does not provide a growth advantage (61). Our results suggest that nutrient availability does not generally affect *in vitro* biofilm formation, but rather does so in a strain-dependent manner. The isolation source is further weakly correlated with their preference of forming stronger biofilms *in vitro* under either nutrient-poor or –rich conditions. Thus, of the isolates changing their biofilm type, fresh produce isolates preferentially formed stronger biofilm under nutrient-poor conditions, while water isolates preferentially formed stronger biofilm under nutrient-rich conditions. The reason for this difference remains elusive and we could not find any previous study comparing *E. coli* isolates from these environmental reservoirs. However, differences in *in vitro* biofilm formation related to nutrient availability and isolation source have been shown for example in avian pathogenic and faecal commensal *E. coli* (62).

Correlation between *E. coli* phylogroups and different phenotypes have previously been reported (63). Consequently, *E. coli* isolates from mammalian hosts are dominated by phylogroup A and B2 while crop hosts are dominated by phylogroup B1. Phylogroup B1 strains are characterised by traits enabling plant colonization. Interestingly, in the present study isolates from phylogroup B1 showed significantly higher average *in vitro* biofilm formation than phylogroup A strains. This difference was, however, observed in both poor and rich media, and does not provide a specific advantage in the resource-limited phyllosphere. Indeed, no such advantage could be detected when testing the strains on gnotobiotic plants. Possibly, the lack of bacterial competition during leaf colonization can explain this lack of signal (64).

From a food safety perspective, the persistence of multidrug-resistant *E. coli* in the phyllosphere of fresh produce is of great importance, as the ingestion of AR bacteria is of concern even when they are not pathogenic. Notably, ten of eleven *E. coli* isolates screened *in planta* had a MAR index exceeding the cut-off of 0.2, which has been widely used to indicate origin from a high-risk source with frequent antibiotic usage (65, 66), and is more common in clinical than environmental isolates (67). Although *in vitro* biofilm formation capacity did not correlate with colonization and survival in the phyllosphere of gnotobiotic lamb’s lettuce, all investigated strains were able to persist and survive on the plants for at least 21 days and did so, more importantly, at unchanged concentrations. The majority of strains maintained stable bacterial loads or could even increase in population density over time. Our results, although stemming from a gnotobiotic system, suggest that an *E. coli* contamination event as long as three weeks prior to harvest might be sufficient to result in persisting populations of *E. coli* in the phyllosphere of fresh produce at harvest. This is in accordance with previous results demonstrating (non-pathogenic and pathogenic) *E. coli* persistence on plant leaves for several weeks under conditions challenging for bacterial survival such as climate-controlled greenhouse (68) or in the open field (69). A comprehensive review on the fate of *E. coli* O157:H7 in leafy greens summarizes 65 studies conducted under such more challenging conditions compared to a gnotobiotic system (70). The reported studies focusing on primary production differ greatly in type of leafy green investigated, initial inoculum (2 – 8 log CFU per mL), observation timeframe (from roughly 4 to 54 days), and detection method (direct plating, enrichment, most probable number (MPN), or qPCR). Keeping in mind possible behavioural differences between non-pathogenic and pathogenic *E. coli*, they all reported declining populations over time (despite long-term detection of the inoculated bacteria in some cases, for up to 28 days), in contrast to our findings of stable or increasing populations. Regarding die-off kinetics, a modelling study by Seidu and colleagues (71) concluded that a biphasic die-off model rather than a log-linear model best described die-off of *E. coli* (not specifically pathogenic strains) on lettuce and cabbage. Of note, quantitative culture-dependent methods are much less sensitive than molecular methods (especially when damaged or dead cells are excluded in molecular methods as in the present study), consequently studies relying on quantitative culture-based strategies usually report on shorter persistence periods. For example, a study by Wood and colleagues investigating survival of antibiotic-resistant *E. coli* inoculated by irrigation onto spinach under commercial production conditions, reported persistence periods in the spinach phyllosphere of less than one week with an exponential decline (72). A study by Schwarz *et al.* showed comparable rapid decay of *E. coli* and *Salmonella enterica* on leaves and spikelets of wheat plants (73).

Interestingly, we did not record clearly increased survival of *in vitro* biofilm-forming strains on leaves, in contrast with the increased bacterial survival within aggregates on leaf surfaces observed in host-adapted epiphytes such as *Pseudomonas syringae* (74). Strikingly, we found that the observed population densities did not follow distributions that could be explained by simple normal, or log-normal distributions as they are normally found in experiments under laboratory conditions (75, 76). Instead, they can be explained best by a mixture of normal distributions. The reasons for this phenomenon is currently unclear.

In summary, this study highlights the complexity of biofilm formation in environmental AR *E. coli* strains. While nutrient availability, isolation source, and phylogroups influence biofilm formation *in vitro*, *in vivo* validation is essential for predicting potential health risks. Despite correlations between *in vitro* biofilm formation and certain traits, no clear advantage was observed in plant colonization. However, all tested strains persisted on lamb’s lettuce, posing a potential food safety concern through unwanted ingestion of AR bacteria. Interestingly, biofilm formation did not consistently enhance survival on leaves, suggesting a more intricate interplay between bacteria and their environment. Further research is needed to understand *E. coli* population dynamics and distribution patterns on plant surfaces. These findings underscore the need for comprehensive approaches to studying biofilm dynamics and their implications for food safety.

## Acknowledgements

We thank Roger Marti for providing expertise on *in vitro* biofilm formation assays and Susanna Lattmann for assistance with laboratory work. We acknowledge Denise Klöti (Agroscope) for providing fluted filter paper and advice on seed sowing and sprouting. This work was part of the Agroscope Research Programme “Reduction and Dynamics of Antibiotic-resistant and Persistent Microorganisms along Food Chains (REDYMO)” and the National Research Program “Antimicrobial Resistance” (NRP 72, grant number 407240_167068) of the Swiss National Science Foundation.

## Code and data availability

Images and raw data are available in Zenodo (77). Scripts and analyses are available in the Github repository: https://github.com/relab-fuberlin/schlechter_biofilm_2024.

## References

1. Su LJ, Arab L. 2006. Salad and raw vegetable consumption and nutritional status in the adult US population: results from the Third National Health and Nutrition Examination Survey. J Am Diet Assoc 106:1394–404.

2. Callejón RM, Rodríguez-Naranjo MI, Ubeda C, Hornedo-Ortega R, Garcia-Parrilla MC, Troncoso AM. 2015. Reported foodborne outbreaks due to fresh produce in the United States and European Union: trends and causes. Foodborne Pathog Dis 12:32–38.

3. Drissner D, Gekenidis M-T. 2023. Safety of Food and Beverages: Fruits and Vegetables. Reference Module in Food Science 1:10–19.

4. Machado-Moreira B, Richards K, Brennan F, Abram F, Burgess CM. 2019. Microbial contamination of fresh produce: what, where, and how? Compr Rev Food Sci Food Saf 18:1727–1750.

5. Rolain J-M. 2013. Food and human gut as reservoirs of transferable antibiotic resistance encoding genes. Front Microbiol 4.

6. Bailey JK, Pinyon JL, Anantham S, Hall RM. 2010. Commensal *Escherichia coli* of healthy humans: a reservoir for antibiotic-resistance determinants. J Med Microbiol 59:1331–1339.

7. Larsson D, Flach C-F. 2022. Antibiotic resistance in the environment. Nature Reviews Microbiology 20:257–269.

8. Blaak H, van Hoek AH, Veenman C, van Leeuwen AED, Lynch G, van Overbeek WM, de Roda Husman AM. 2014. Extended spectrum β-lactamase- and constitutively AmpC-producing *Enterobacteriaceae* on fresh produce and in the agricultural environment. Int J Food Microbiol 168:8–16.

9. Blau K, Bettermann A, Jechalke S, Fornefeld E, Vanrobaeys Y, Stalder T, Top EM, Smalla K. 2018. The Transferable Resistome of Produce. MBio 9:e01300–18.

10. Kim M-C, Cha M-H, Ryu J-G, Woo G-J. 2017. Characterization of vancomycin-resistant *Enterococcus faecalis* and *Enterococcus faecium* isolated from fresh produces and human fecal samples. Foodborne Pathog Dis 14:195–201.

11. Kläui A, Bütikofer U, Naskova J, Wagner E, Marti E. 2024. Fresh produce as a reservoir of antimicrobial resistance genes: A case study of Switzerland. Sci Total Environ 907:167671.

12. Nüesch-Inderbinen M, Zurfluh K, Peterhans S, Hächler H, Stephan R. 2015. Assessment of the prevalence of extended-spectrum beta-lactamase-producing *Enterobacteriaceae* in ready-to-eat salads, fresh-cut fruit, and sprouts from the Swiss market. J Food Prot 78:1178–81.

13. Reid CJ, Blau K, Jechalke S, Smalla K, Djordjevic SP. 2020. Whole Genome Sequencing of *Escherichia coli* From Store-Bought Produce. Front Microbiol 10:3050.

14. Vital PG, Caballes MBD, Rivera WL. 2017. Antimicrobial resistance in *Escherichia coli* and *Salmonella* spp. isolates from fresh produce and the impact to food safety. J Environ Sci Health, B 52:683–689.

15. who.int [Internet]. World Health Organization (WHO): Antibiotic resistance - fact sheet; c2020. https://www.who.int/en/news-room/fact-sheets/detail/antibiotic-resistance. Accessed February 2023.

16. Da Silva SF, Reis IB, Monteiro MG, Dias VC, Machado ABF, Da Silva VL, Diniz CG. 2021. Influence of human eating habits on antimicrobial resistance phenomenon: Aspects of clinical resistome of gut microbiota in omnivores, ovolactovegetarians, and strict vegetarians. Antibiotics 10:276.

17. O’Flaherty E, Solimini AG, Pantanella F, De Giusti M, Cummins E. 2018. Human exposure to antibiotic resistant-*Escherichia coli* through irrigated lettuce. Environ Int 122:270–280.

18. Schlechter RO, Miebach M, Remus-Emsermann MNP. 2019. Driving factors of epiphytic bacterial communities: A review. J Adv Res 19:57–65.

19. Leveau JHJ, Lindow SE. 2001. Appetite of an epiphyte: Quantitative monitoring of bacterial sugar consumption in the phyllosphere. PNAS 98:3446–3453.

20. Monier J-M, Lindow SE. 2004. Frequency, Size, and Localization of Bacterial Aggregates on Bean Leaf Surfaces. Appl Environ Microbiol 70:346–355.

21. Muhammad MH, Idris AL, Fan X, Guo Y, Yu Y, Jin X, Qiu J, Guan X, Huang T. 2020. Beyond Risk: Bacterial Biofilms and Their Regulating Approaches. Front Microbiol 11:928.

22. de Carvalho CCCR. 2017. Biofilms: Microbial Strategies for Surviving UV Exposure, p 233-239. In Ahmad S (ed), Ultraviolet Light in Human Health, Diseases and Environment; Advances in Experimental Medicine and Biology, vol 996. Springer, Cham.

23. Allison DG. 2003. The biofilm matrix. Biofouling 19:139–150.

24. Danhorn T, Fuqua C. 2007. Biofilm formation by plant-associated bacteria. Annu Rev Microbiol 61:401–422.

25. Gil MI, Selma MV, López-Gálvez F, Allende A. 2009. Fresh-cut product sanitation and wash water disinfection: problems and solutions. Int J Food Microbiol 134:37–45.

26. Yaron S, Römling U. 2014. Biofilm formation by enteric pathogens and its role in plant colonization and persistence. Microb Biotechnol 7:496–516.

27. Artés F, Gómez P, Aguayo E, Escalona V, Artés-Hernández F. 2009. Sustainable sanitation techniques for keeping quality and safety of fresh-cut plant commodities. Postharvest Biol Technol 51:287–296.

28. Modesti M, Macaluso M, Taglieri I, Bellincontro A, Sanmartin C. 2021. Ozone and Bioactive Compounds in Grapes and Wine. Foods 10:2934.

29. McBain AJ. 2009. *In vitro* biofilm models: an overview. Adv Appl Microbiol 69:99–132.

30. Lapidot A, Romling U, Yaron S. 2006. Biofilm formation and the survival of *Salmonella* Typhimurium on parsley. Int J Food Microbiol 109:229–233.

31. Lapidot A, Yaron S. 2009. Transfer of *Salmonella enterica* serovar Typhimurium from contaminated irrigation water to parsley is dependent on curli and cellulose, the biofilm matrix components. J Food Prot 72:618–23.

32. Cevallos-Cevallos JM, Gu G, Danyluk MD, van Bruggen AH. 2012. Adhesion and splash dispersal of *Salmonella enterica* Typhimurium on tomato leaflets: effects of rdar morphotype and trichome density. Int J Food Microbiol 160:58–64.

33. Cowles KN, Willis DK, Engel TN, Jones JB, Barak JD. 2016. Diguanylate Cyclases AdrA and STM1987 Regulate *Salmonella enterica* Exopolysaccharide Production during Plant Colonization in an Environment-Dependent Manner. Appl Environ Microbiol 82:1237–1248.

34. Niemira BA, Cooke PH. 2010. *Escherichia coli* O157:H7 Biofilm Formation on Romaine Lettuce and Spinach Leaf Surfaces Reduces Efficacy of Irradiation and Sodium Hypochlorite Washes. J Food Sci 75:M270–M277.

35. Klug TV, Novello J, Laranja DC, Aguirre TAS, de Oliveira Rios A, Tondo EC, Santos RPd, Bender RJ. 2017. Effect of Tannin Extracts on Biofilms and Attachment of *Escherichia coli* on Lettuce Leaves. Food and Bioprocess Technology 10:275–283.

36. Zhang Y, Huang HH, Ma LZ, Masuda Y, Honjoh KI, Miyamoto T. 2022. Inactivation of mixed Escherichia coli O157:H7 biofilms on lettuce by bacteriophage in combination with slightly acidic hypochlorous water (SAHW) and mild heat treatment. Food Microbiol 104:104010.

37. Ahrens S. 2022. Ertrag durch den Anbau von verschiedenen Salatsorten in Deutschland von 2019 bis 2021. https://de.statista.com/statistik/daten/studie/639108/umfrage/ertragsmenge-von-salatsorten-in-deutschland. Accessed February 2023.

38. Bundesamt für Landwirtschaft BLW. 2016. Marktbericht Früchte und Gemüse - Nüsslisalat liegt im Trend.

39. Ettlin I. 2022. Nüsslisalat – der Gesundheits-Champion, p *In* BauernZeitung. Schweizer Agrarmedien AG, Münchenbuchsee, Switzerland. https://www.bauernzeitung.ch/artikel/pflanzen/nuesslisalat-der-gesundheits-champion-448102#:~:text=Beliebter%20Wintersalat&text=F%C3%BCr%20die%20Gem%C3%BCseproduzenten%20und%20%2Dproduzentinnen,dem%20Kopfsalat%20auf%20Platz%202.

40. Gekenidis MT, Schoner U, von Ah U, Schmelcher M, Walsh F, Drissner D. 2018. Tracing back multidrug-resistant bacteria in fresh herb production: from chive to source through the irrigation water chain. FEMS Microbiol Ecol 94:fiy149.

41. Clermont O, Christenson JK, Denamur E, Gordon DM. 2013. The Clermont *Escherichia coli* phylo-typing method revisited: improvement of specificity and detection of new phylo-groups. Environ Microbiol Rep 5:58–65.

42. Reisner A, Krogfelt KA, Klein BM, Zechner EL, Molin S. 2006. In vitro biofilm formation of commensal and pathogenic *Escherichia coli* strains: impact of environmental and genetic factors. J Bacteriol 188:3572–3581.

43. Reisner A, Krogfelt KA, Klein BM, Zechner EL, Molin S. 2006. In Vitro Biofilm Formation of Commensal and Pathogenic *Escherichia coli* Strains: Impact of Environmental and Genetic Factors. J Bacteriol 188:3572–3581.

44. Marti R, Schmid M, Kulli S, Schneeberger K, Naskova J, Knøchel S, Ahrens CH, Hummerjohann J. 2017. Biofilm formation potential of heat-resistant *Escherichia coli* dairy isolates and the complete genome of multidrug-resistant, heat-resistant strain FAM21845. Appl Environ Microbiol 83:e00628–17.

45. Peng S, Stephan R, Hummerjohann J, Blanco J, Zweifel C. 2012. In vitro characterization of Shiga toxin-producing and generic *Escherichia coli* in respect of cheese-production relevant stresses. J Food Saf Food Qual 63:136–141.

46. Stepanović S, Vuković D, Hola V, di Bonaventura G, Djukić S, Ćirković I, Ruzicka F. 2007. Quantification of biofilm in microtiter plates: overview of testing conditions and practical recommendations for assessment of biofilm production by staphylococci. APMIS 115:891–899.

47. Kremer JM, Sohrabi R, Paasch BC, Rhodes D, Thireault C, Schulze-Lefert P, Tiedje JM, He SY. 2021. Peat-based gnotobiotic plant growth systems for Arabidopsis microbiome research. Nat Protoc 16:2450–2470.

48. Ma KW, Ordon J, Schulze-Lefert P. 2022. Gnotobiotic plant systems for reconstitution and functional studies of the root microbiota. Current protocols 2:e362.

49. Miebach M, Schlechter RO, Clemens J, Jameson PE, Remus-Emsermann MN. 2020. Litterbox—a gnotobiotic zeolite-clay system to investigate Arabidopsis–microbe interactions. Microorganisms 8:464.

50. Vorholt JA, Vogel C, Carlström CI, Müller DB. 2017. Establishing Causality: Opportunities of Synthetic Communities for Plant Microbiome Research. Cell Host & Microbe 22:142–155.

51. Elizaquível P, Aznar R, Sánchez G. 2014. Recent developments in the use of viability dyes and quantitative PCR in the food microbiology field. J Appl Microbiol 116:1–13.

52. Clifford RJ, Milillo M, Prestwood J, Quintero R, Zurawski DV, Kwak YI, Waterman PE, Lesho EP, Mc Gann P. 2012. Detection of bacterial 16S rRNA and identification of four clinically important bacteria by real-time PCR. PLoS One 7:e48558.

53. R Core Team. 2021. R: A language and environment for statistical computing. R Foundation for Statistical Computing, Vienna, Austria.

54. Wickham H, Averick M, Bryan J, Chang W, McGowan LDA, François R, Grolemund G, Hayes A, Henry L, Hester J. 2019. Welcome to the Tidyverse. J Open Source Softw 4:1686.

55. Pinheiro J, Bates D, DebRoy S, Sarkar D, R Core Team. 2021. nlme: Linear and Nonlinear Mixed Effects Models.

56. Lenth RV. 2022. Emmeans: Estimated Marginal Means, aka Least-Squares Means, Version 1.8. 1–1.

57. Benaglia T, Chauveau D, Hunter DR, Young DS. 2009. mixtools: An R Package for Analyzing Mixture Models. J Stat Softw 32:1–29.

58. Magiorakos AP, Srinivasan A, Carey RB, Carmeli Y, Falagas ME, Giske CG, Harbarth S, Hindler JF, Kahlmeter G, Olsson-Liljequist B. 2012. Multidrug-resistant, extensively drug-resistant and pandrug-resistant bacteria: an international expert proposal for interim standard definitions for acquired resistance. Clin Microbiol Infect 18:268–281.

59. Roberts AEL, Kragh KN, Bjarnsholt T, Diggle SP. 2015. The Limitations of *In Vitro* Experimentation in Understanding Biofilms and Chronic Infection. J Mol Biol 427:3646–3661.

60. Ferreira FA, Souza RR, Bonelli RR, Américo MA, Fracalanzza SEL, Figueiredo AMS. 2012. Comparison of in vitro and *in vivo* systems to study ica-independent *Staphylococcus aureus* biofilms. J Microbiol Methods 88:393–398.

61. Stanley NR, Lazazzera BA. 2004. Environmental signals and regulatory pathways that influence biofilm formation. Mol Microbiol 52:917–924.

62. Skyberg JA, Siek KE, Doetkott C, Nolan LK. 2007. Biofilm formation by avian *Escherichia coli* in relation to media, source and phylogeny. J Appl Microbiol 102:548–554.

63. Méric G, Kemsley EK, Falush D, Saggers EJ, Lucchini S. 2013. Phylogenetic distribution of traits associated with plant colonization in *Escherichia coli*. Environ Microbiol 15:487–501.

64. Schlechter RO, Remus-Emsermann MN. 2023. Bacterial community complexity in the phyllosphere penalises specialists over generalists. bioRxiv doi:10.1101/2023.11.08.566251.

65. Krumperman PH. 1983. Multiple antibiotic resistance indexing of *Escherichia coli* to identify high-risk sources of fecal contamination of foods. Appl Environ Microbiol 46:165–170.

66. Titilawo Y, Sibanda T, Obi L, Okoh A. 2015. Multiple antibiotic resistance indexing of *Escherichia coli* to identify high-risk sources of faecal contamination of water. Environ Sci Pollut Res 22:10969–10980.

67. Beattie RE, Bakke E, Konopek N, Thill R, Munson E, Hristova KR. 2020. Antimicrobial resistance traits of *Escherichia coli* isolated from dairy manure and freshwater ecosystems are similar to one another but differ from associated clinical isolates. Microorganisms 8:747.

68. Gekenidis M-T, Rigotti S, Hummerjohann J, Walsh F, Drissner D. 2020. Long-Term Persistence of *bla*_CTX-M-15_ in Soil and Lettuce after Introducing Extended-Spectrum β-Lactamase (ESBL)-Producing *Escherichia coli* via Manure or Water. Microorganisms 8:1646.

69. Mark Ibekwe A, Grieve CM, Papiernik SK, Yang CH. 2009. Persistence of *Escherichia coli* O157:H7 on the rhizosphere and phyllosphere of lettuce. Lett Appl Microbiol 49:784–790.

70. Owade JO, Bergholz TM, Mitchell J. 2024. A review of conditions influencing fate of Shiga toxin-producing *Escherichia coli* O157:H7 in leafy greens. Compr Rev Food Sci Food Saf 23:e70013.

71. Seidu R, Sjølander I, Abubakari A, Amoah D, Larbi JA, Stenström TA. 2013. Modeling the die-off of *E. coli* and *Ascaris* in wastewater-irrigated vegetables: implications for microbial health risk reduction associated with irrigation cessation. Water Sci Technol 68:1013–1021.

72. Wood JD, Bezanson GS, Gordon RJ, Jamieson R. 2010. Population dynamics of *Escherichia coli* inoculated by irrigation into the phyllosphere of spinach grown under commercial production conditions. Int J Food Microbiol 143:198–204.

73. Schwarz K, Sidhu JPS, Pritchard D, Li Y, Toze S. 2014. Decay of *Salmonella enterica*, *Escherichia coli* and bacteriophage MS2 on the phyllosphere and stored grains of wheat (*Triticum aestivum*). Lett Appl Microbiol 58:16–24.

74. Monier J-M, Lindow SE. 2003. Differential survival of solitary and aggregated bacterial cells promotes aggregate formation on leaf surfaces. PNAS 100:15977–15982.

75. Remus-Emsermann MNP, Aicher D, Pelludat C, Gisler P, Drissner D. 2021. Conjugation Dynamics of Self-Transmissible and Mobilisable Plasmids into *E. coli* O157:H7 on *Arabidopsis thaliana* Rosettes. Antibiotics 10:928.

76. Schlechter RO, Kear EJ, Bernach M, Remus DM, Remus-Emsermann MNP. 2023. Metabolic resource overlap impacts competition among phyllosphere bacteria. ISME J 17:1445–1454.

77. Schlechter R, Marti E, Remus-Emsermann M, Drissner D, Gekenidis M-T. 2024. Data set “Correlation of in vitro biofilm formation capacity with persistence of antibiotic-resistant Escherichia coli on fresh leafy produce” [Data set] doi:10.5281/zenodo.13838842.

