## Supplemental Files for "Correlation of *in vitro* biofilm formation capacity with persistence of antibiotic-resistant *Escherichia coli* on gnotobiotic lamb’s lettuce"

**Correlation of *in vitro* biofilm formation capacity with persistence of antibiotic-resistant *Escherichia coli* on fresh leafy produce**

Rudolf O. Schlechter^a^, Elisabet Marti^b^, Mitja N. P. Remus-Emsermann^a^, David Drissner^c^, and Maria-Theresia Gekenidis^d^*

^a^ Institute of Microbiology, Department of Biology, Chemistry, Pharmacy, Freie Universität Berlin, 14195 Berlin, Germany

^b^ Research Group Microbiological Food Safety, Agroscope, 3003 Liebefeld, Switzerland

^c^ Department of Life Sciences, Albstadt-Sigmaringen University, 72488 Sigmaringen, Germany

^d^ Research Division Food Microbial Systems, Agroscope, 8820 Wädenswil, Switzerland

**Supplemental Table**

**Table S1.** Change in biofilm type category of isolated environmental *E. coli* strains relative to the assignment in ABTCAA medium. BF: biofilm formation.

| ***Isolation source*** | **Total # of strains** | **Increased BF in** **LB-NaCl(0)** | **Decreased BF in LB-NaCl(0)** | **Unchanged BF** |
| --- | --- | --- | --- | --- |
| **Fresh produce** | **35** | 9 (25.7%) | 17 (48.6%) | 9 (25.7%) |
| **Soil** | **20** | 4 (20.0 %) | 4 (20.0%) | 12 (60.0%) |
| **Water** | **117** | 40 (34.2%) | 20 (17.1%) | 57 (48.7%) |
| **Greenhouse (sprinkler)** | **2** | 1 (50.0%) | 0 (0.0 %) | 1 (50.0%) |
| **Total** | **174** | 54 | 41 | 79 |

Percentages (%) are in relation to the total number of strains from each isolation source.

**Supplemental Figures**

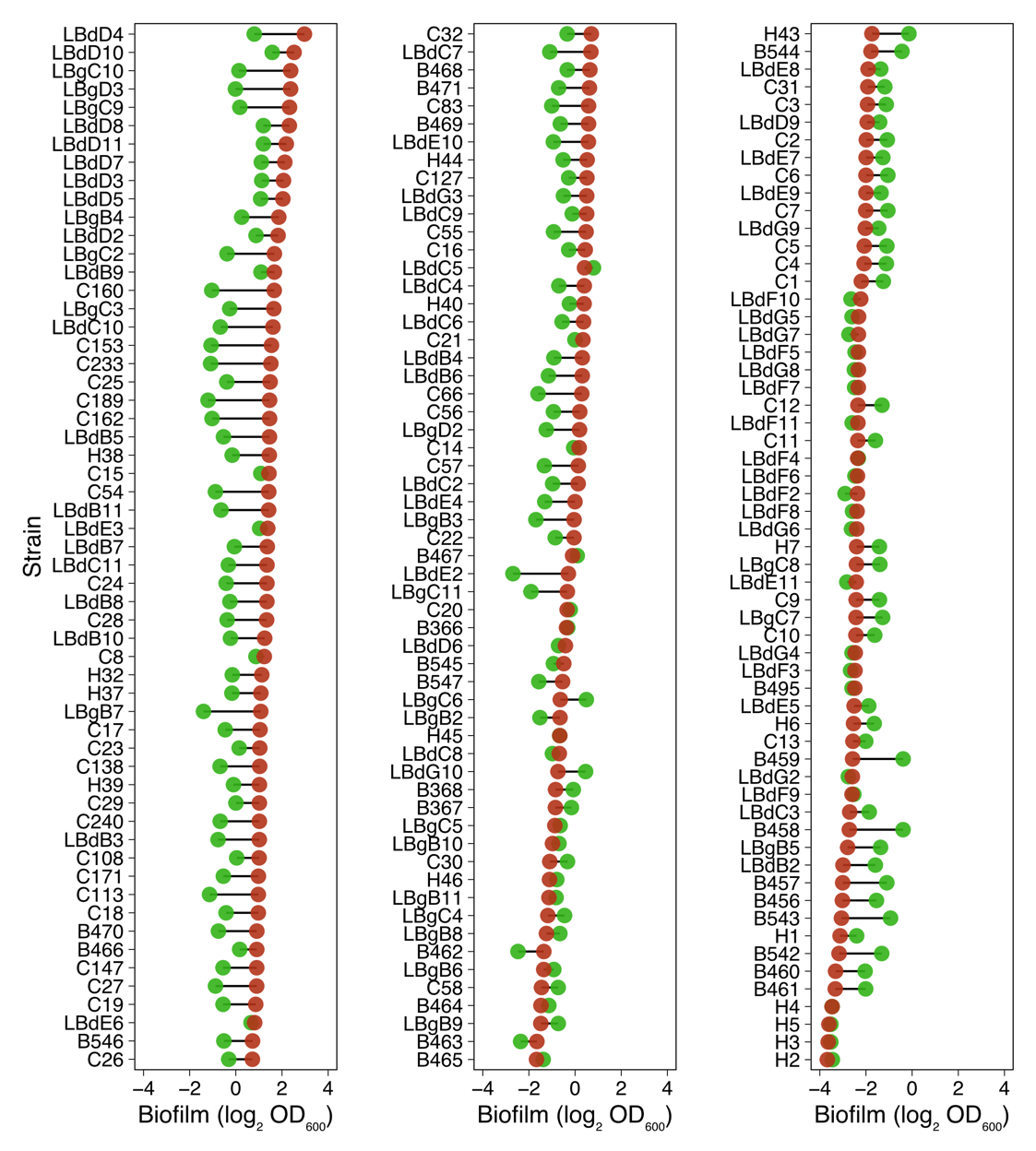

**Figure S1.** Differences in biofilm capacity of every tested strain in ABTCAA (green) or LB-NaCl(0) (red). Values indicate the average log_2_-transformed OD_600_ values from crystal violet assays.

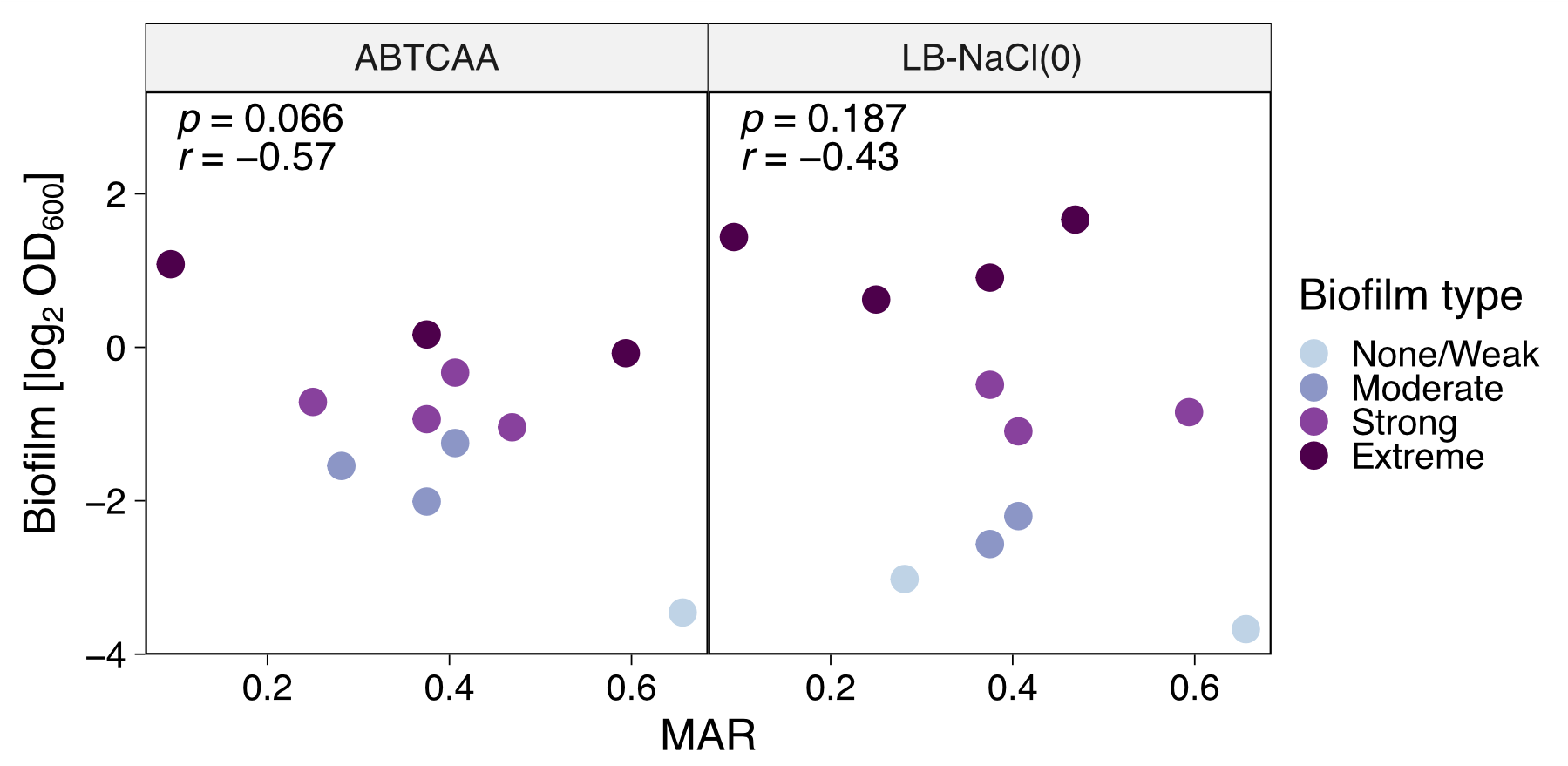

**Figure S2.** Correlation between biofilm formation and antibiotic resistance *in vitro*. MAR, multiple antibiotic resistance index.

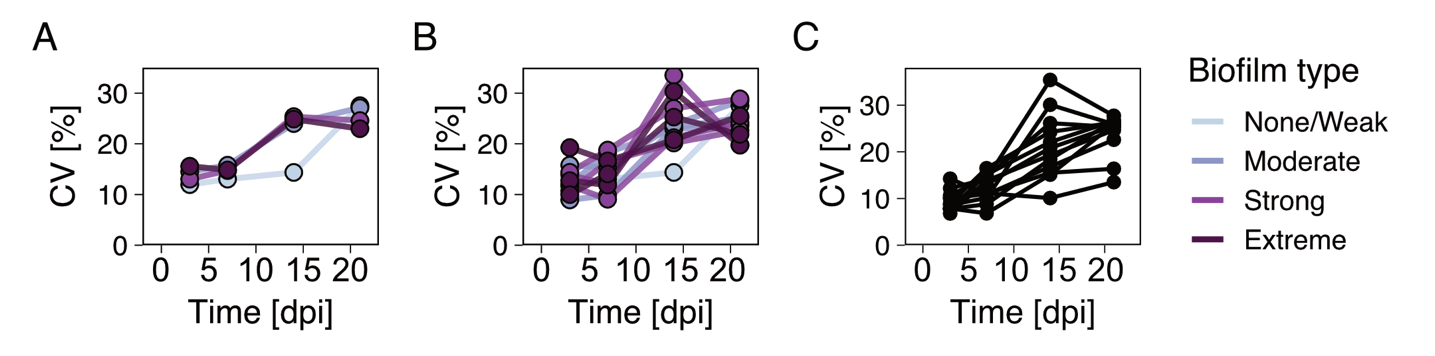

**Figure S3.** Coefficient of variation (CV) of qPCR data in the phyllosphere. The extent of the variability in bacterial abundance on leaves was calculated as a coefficient of variation (%). Change in coefficient of variation over time of *E. coli* populations on *V. locusta*, grouped by biofilm category (A), as individual strain (B), or independent experiments (C). dpi, days post inoculation.

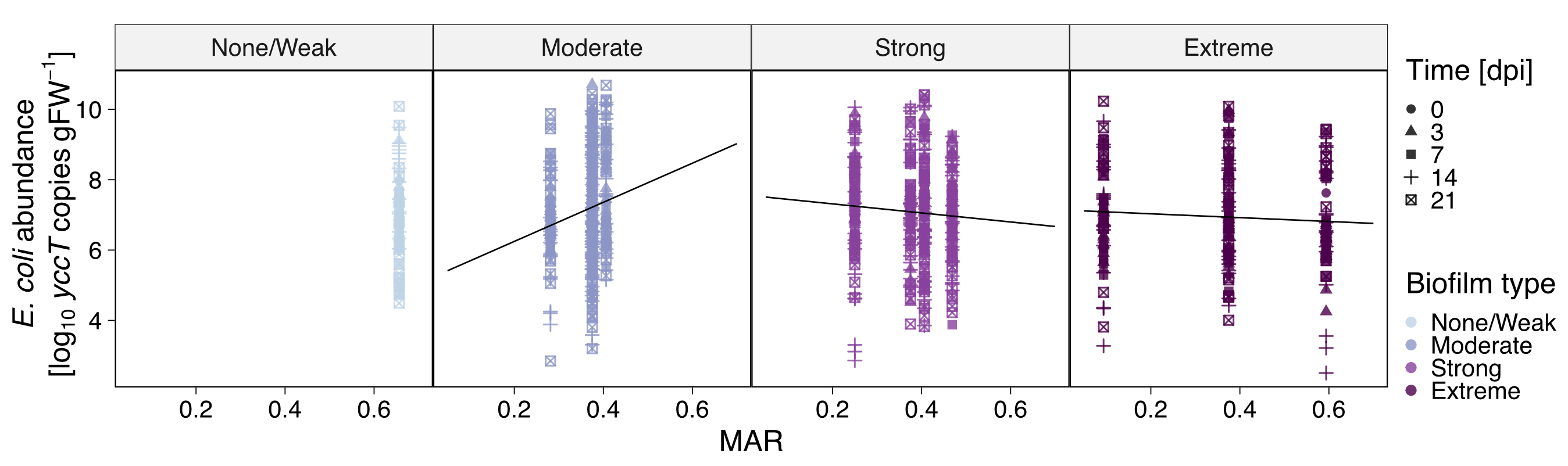

**Figure S4.** Correlation between biofilm-forming *E. coli* and antibiotic resistance *in planta*. gFW, gram fresh weight; dpi, days post inoculation; MAR, multiple antibiotic resistance index.
